# Unbiased Assessment of H-STS cells as high-fidelity models for gastro-enteropancreatic neuroendocrine tumor drug mechanism of action analysis

**DOI:** 10.1101/677435

**Authors:** Mariano J. Alvarez, Pengrong Yan, Mary L. Alpaugh, Michaela Bowden, Ewa Sicinska, Chensheng W. Zhou, Charles Karan, Ronald B. Realubit, Prabhjot S. Mundi, Adina Grunn, Dirk Jäger, John A. Chabot, Antonio T. Fojo, Paul E. Oberstein, Hanina Hibshoosh, Jeffrey W. Milsom, Matthew H. Kulke, Massimo Loda, Gabriela Chiosis, Diane L. Reidy-Lagunes, Andrea Califano

## Abstract

Quantitative metrics to objectively assess the fidelity of cancer models, such as cell lines, organoids, or patient-derived xenografts (PDXs), remain elusive, with histological criteria or the presence of specific mutations often used as driving principles. We show that molecular criteria, based on the regulatory proteins responsible for maintaining transcriptional cell state and its regulatory network, are effective in identifying models that can recapitulate drug mechanism of action and drug sensitivity, independent of histological consideration.

## Introduction

What constitutes an appropriate model to study the biology of a specific tumor remains the matter of considerable debate, with histology-based or mutational criteria being frequently used as *ad hoc* criteria for model-fidelity. Unfortunately, the latter can be misleading and negatively bias model selection. For instance, to highlight potential pitfalls of mutation-based fidelity assessment, we recently evaluated the suitability of 16 neuroblastoma cell lines harboring MYCN amplifications, as high-fidelity models for the study of the MYCN-dependent (MYCNA) subtype of this tumor (1). Surprisingly, only four of these effectively recapitulated key tumor dependencies and associated mechanisms that were shown to be conserved across virtually all MYCN-subtype patients, in two independent cohorts. Indeed, focusing on individual mutations— albeit important ones, associated with tumor etiology—will inevitably ignore the effect of a large complement of additional, model-specific genetic and epigenetic events.

Currently, effective assessment of tumor-model fidelity is best performed *a posteriori*, for instance, by determining whether tumor dependencies identified in a tumor model (e.g., a cell line or organoid) are recapitulated *in vivo* in a Patient Derived Xenograft (PDX) or, even better, in patients (e.g., via a clinical trial). However, these approaches are obviously both highly inefficient and time-consuming and would greatly benefit from the availability of methodologies capable of quantitatively assess model fidelity *a priori*. Equally important, model-fidelity may not be assessed generically but only in the context of the specific questions being asked. For instance, a high-fidelity model to identify genes that are essential to tumor viability (tumor dependencies) may not be equally appropriate to identify drug sensitivity biomarkers.

Taken together, these observations suggest that, while critically needed, objective metrics to assess model-fidelity are still elusive. This motivated us to develop a quantitative, molecular-level framework (*OncoMatch*) to assess the fidelity of a given tumor model in the context of the specific biological question being asked. In (2), we proposed addressing this challenge by integrating two independent metrics: (a) conservation of regulatory networks inferred exclusively from patient-derived samples in the tumor-model and (b) overlap of patient-specific Master Regulator (MR) proteins— i.e., proteins representing the mechanistic determinants of the transcriptional state associated with the phenotype of interest (3)—with tumor-model-specific MRs. The latter are identified based on the enrichment of their positively regulated and repressed targets in the phenotype-specific transcriptional signature, using the Virtual Inference of Protein activity by Enriched Regulon (VIPER) algorithm, which has been extensively validated (4). Such an unbiased criterion would be especially valuable for tumors that lack effective *in vitro* or *in vivo* models (e.g. carcinoids or prostate cancer), or even when an optimal model must be chosen among several available ones, such as in neuroblastoma.

While the rationale for the proposed methodology is biologically sound, the original manuscript did not provide experimental data supporting it. As a result, the two cell lines prioritized by OncoMatch as high-fidelity Gastro-EnteroPancreatic NeuroEndocrine Tumors (GEP-NETs) models (2) (H-STS and KRJ-1) have recently come under scrutiny. Specifically, they were reported as derived from Epstein-Barr Virus (EBV)-immortalized lymphoblastoid cells (5) rather than from *bona fide* GEP-NET tumor cells, thus violating a key tenet of traditional model-fidelity assessment.

In this manuscript, we confirm that H-STS and KRJ-1 cells represent high-fidelity models for the assessment of drug mechanism of action and drug sensitivity in GEP-NETs, by leveraging an extensive, novel set of drug-perturbation assays in primary cells and explants derived from GEP-NET patients. This suggests that quantitatively motivated assessments, such as OncoMatch may effectively complement and extend histology and mutational-based criteria in assessing tumor model fidelity.

## Results

OncoMatch analysis comprises the following three steps: (a) assessing whether a regulatory model elucidated exclusively from patient-derived tumor profiles recapitulates molecular interactions in the proposed tumor model (b) assessing the overlap between patient-derived MR proteins with MR proteins inferred from the candidate model and (c) integrating the p-values generated by the two metrics using Stouffer’s method, see (2) for further details on the approach.

For the GEP-NET study, we used OncoMatch to prioritize 923 cell lines as high fidelity models for the study of drug mechanism of action and sensitivity in GEP-NET patients (2). Cell lines included those in the Cancer Cell Line Encyclopedia (CCLE) (6), as well as H-STS and KRJ-1 cells, previously reported as derived from a small-bowel NET patient (7). For VIPER analysis of patient-derived tumors, we used a metastatic progression signature based on the genes differentially expressed between each hepatic metastasis and a set of cluster-matched primary tumors (2). For cell lines, differential expression was computed against P-STS cells, representing a *bona fide* primary small bowel GEP-NET model, as confirmed by mutational analysis (8).

From this comprehensive analysis, H-STS and KRJ-1 emerged as the 4^th^ and 6^th^ highest-fidelity models (Suppl. Fig. 4 in (2)), respectively, and were further selected for their established ability to grow as xenografts and because they were originally reported as GEP-NET patient derived.

Since the identity of these cells was later revised as B-cell-derived (5), we first asked whether other B-cell-derived lines in CCLE would also emerge as high-fidelity models for GEP-NET studies or whether H-STS and KRJ-1 cells were clear outliers. As shown in Fig. 1, OncoMatch analysis of other normal and transformed B cells consistently identified them as low-fidelity models for GEP-NETs, with a median average rank of 591, across all cell lines, and the top cell line — ST486, an EBV-negative Burkitt’s lymphoma— ranked 22^nd^. These data suggest that H-STS and KRJ-1 constitute, at the very least, highly unusual B-cell-derived lines in terms of their ability to recapitulate the regulatory network and MR activity of 36% (25/69) of metastatic GEP-NET patients, and up to 65% (45/69), when xenografted in immunocompromised mice (Fig. 2 in (2)). Consistently, use of these cells as GEP-NET models was limited to patient samples with high OncoMatch-based priority (*p <* 10^*−*10^, Bonferroni Corrected (BC)).

**Fig. 1.**
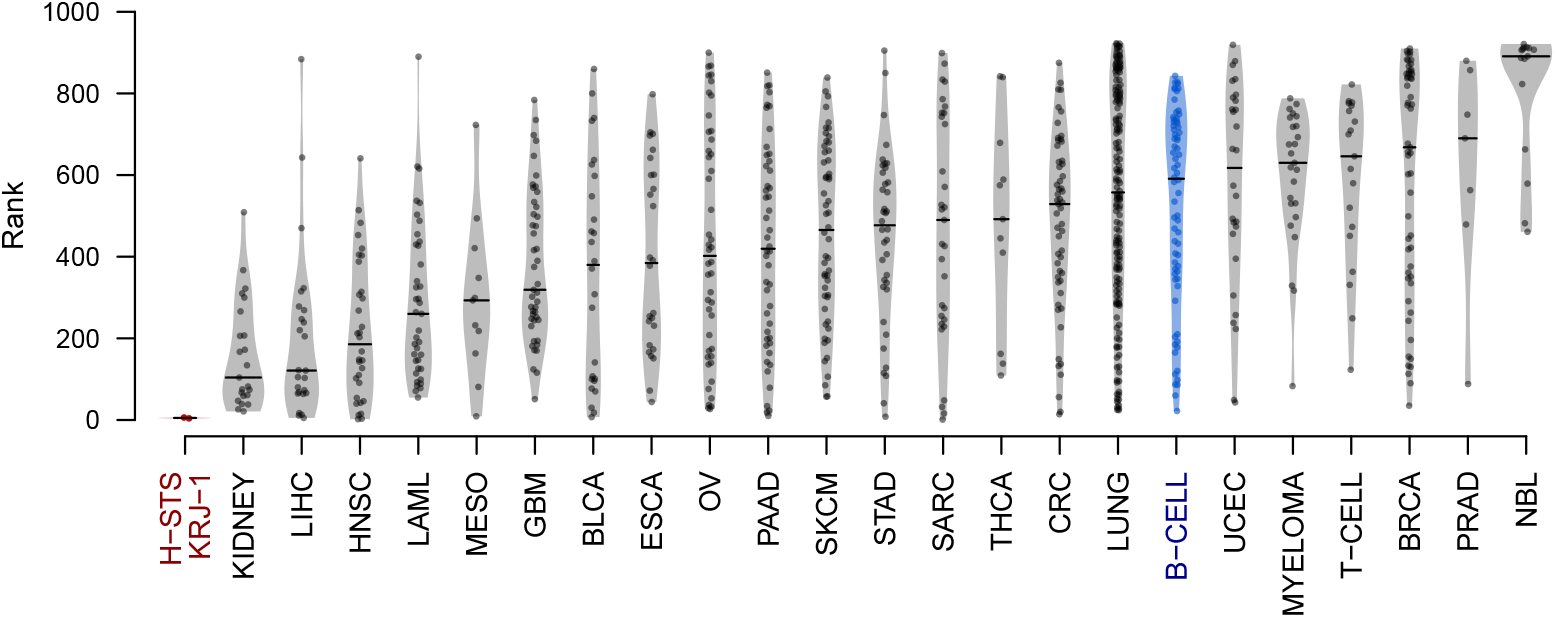
GEP-NET model score for 923 cell lines. Distribution of the model score, computed by integrating the GEP-NET interactome network-score and the number of MR-matched metastases, as described in (2), for the H-STS and KRJ-1 cells (red), and 921 cell lines from the Cancer Cell Line Encyclopedia (CCLE) (6), which were grouped by tissue type, including: B-cells (blue), T-cells, kidney, myeloma, liver hepatocellular carcinoma (LIHC), head and neck squamous carcinoma (HNSC), acute myeloid leukemia (LAML), mesothelioma (MESO), glioblastoma (GBM), bladder urothelial carcinoma (BLCA), esophageal carcinoma (ESCA), ovarian carcinoma (OV), pancreas adenocarcinoma (PAAD), skin cutaneous melanoma (SKCM), stomach adenocarcinoma (STAD), sarcoma (SARC), thyroid cancer (THCA), colorectal carcinoma (CRC), uterine corpus endometrial carcinoma (UCEC), breast carcinoma (BRCA), prostate carcinoma (PRAD), neuroblastoma (NBL).

**Fig. 2.**
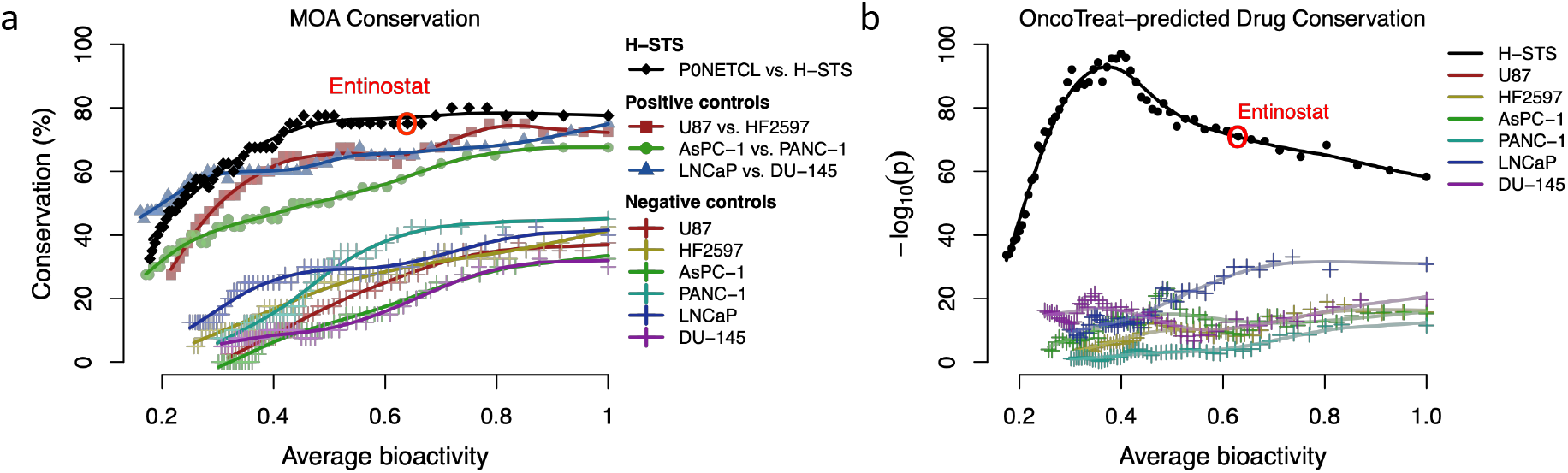
MoA and OncoTreat-prediction conservation. (a) Proportion of drugs showing conserved MoA (*p <* 10^*−*10^, BC), averaged over a running-window of 40 drugs, ranked from lowest to highest bioactivity (see methods), between (*i*) P0NETCL and H-STS cells (solid black diamonds), (*ii*) tumor-matched cell line pairs (positive controls) (colored solid markers) and (*iii*) P0NETCL and six low-fidelity cell line models (negative controls) (colored crosses). (b) Conservation of H-STS and P0NETCL-based OncoTreat-predictions across 69 GEP-NET hepatic metastases, computed by Fisher’s exact test over the same 40-drug running-window basis (solid black circles). Negative control curves were generated by comparing P0NETCL-based OncoTreat predictions to those using perturbational profiles from 6 low-fidelity cell line models (colored crosses).

To further and more objectively evaluate the suitability of these cell lines for OncoTreat analysis of GEP-NETs, as well as to corroborate the published results (2), we report on three novel and distinct analyses made possible by the recent completion of drug screening assays with 126 drugs in primary, rectal-NET patient derived cells (P0NETCL) and with selected, OncoTreat-predicted drugs in GEP-NET-patient-derived explants, cultured and treated in organotypic culture conditions.

Of the drugs originally profiled in H-STS cells (2), 95 were also profiled in P0NETCL cells, supporting direct comparison of Mechanism of Action (MoA) inference based on the overlap of proteins, whose activity was significantly affected by the drug (9), see Methods. Perturbational profiles include RNASeq data generated using the PLATESeq (10) methodology at 24h following cell perturbation with each of the 126 compounds. For each compound, the maximum sublethal concentration (ED_20_) —based on 10-point dose response curves in triplicate, as originally described in (2)— and 1/5^th^ of that concentration were profiled in duplicate.

MoA conservation between P0NETCL and H-STS was highly significant, with 60 of the 95 assessed drugs (63%) showing highly significant MoA similarity at a highly conservative statistical threshold (*p <* 10^*−*10^, BC), suggesting that H-STS cells accurately recapitulate the MoA inferred from *bona fide* GEP-NET cells (Fig. 2a). Indeed, MoA conservation was at least as good as conservation between positive control pairs represented by tumor-type-matched cell lines, for which perturbational profiles were also available. These include U87 and HF2597 (Glioblastoma (GBM)), AsPC-1 and PANC-1 (Pancreas Adenocarcinoma (PDAC)), and LNCaP and DU-145 (Prostate Adenocarcinoma (PRCA)). In particular, MoA similarity for entinostat in P0NETCL and H-STS was highly significant (*p <* 10^*−*11^, BC, see red dot on Fig. 2a). This is relevant, since entinostat was predicted to effectively reverse MR-activity in the largest subset of GEP-NET patients (2) and is currently being tested in a clinical trial. In sharp contrast, MoA conservation between P0NETCL and each of the six cell lines (negative controls) was poor, consistent with their assessment as low-fidelity GEP-NET model by OncoMatch

We then assessed overlap of OncoTreat predictions in 69 GEP-NET hepatic metastases (*p <* 10^*−*5^, BC), using perturbational profiles from either H-STS or P0NETCL cells (see Methods and Fig. 2b). We observed maximum OncoTreat prediction reproducibility at intermediate levels of compound bioactivity, suggesting that drugs with very high bioactivity levels may be more pleiotropic, thus resulting in lower MoA specificity. Consistent with MoA conservation, we observed significant OncoTreat prediction overlap for P0NETCL and H-STS cells (*p <* 10^*−*130^, Fisher’s Exact Test (FET)); odds ratio = 6.7), significantly stronger than the one for P0NETCL cells and the six low-fidelity cell line models (negative controls) (*p* = 0.015, U-test; odds ratio = 2.39 ± 0.29; median ± median absolute deviation). This further confirms OncoMatch predictions of H-STS cells as high-fidelity models for the study of GEP-NET drug activity, based on both overall MoA and OncoTreat prediction conservation.

In an effort to further mitigate technical artifacts associated with primary cultures, we generated perturbational profiles of explants derived from hepatic metastases of GEP-NET patients. These retain the three-dimensional architecture of the tissue and provide more faithful representation of stromal and extracellular matrix contributions. Since explants had to be treated within 24h, their drug sensitivity could not be assessed *a priori* by OncoTreat. As a result, drugs were pre-prioritized based on top OncoTreat predictions at the primary site level, including pancreatic (P-NET), small-bowel (SI-NET), and rectal (RE-NET) (Supplementary Table 1). Each explant was treated with as many drugs as allowed by tissue availability. From 14 explants with sufficient tissue availability to test at least 3 drugs, eight were confirmed as *bona fide* GEP-NET matches, based on transcriptome analysis (Fig. 3), see methods, and were used in subsequent analyses. This allowed assessing experimental reversal of MR-activity following treatment with selected drugs, based on actual (*a posteriori*) OncoTreat predictions using the RNASeq of each explant and using the H-STS perturbational profiles. A conservative statistical significance threshold was used to assess MR-activity reversal (*p <* 10^*−*5^, BC).

**Fig. 3.**
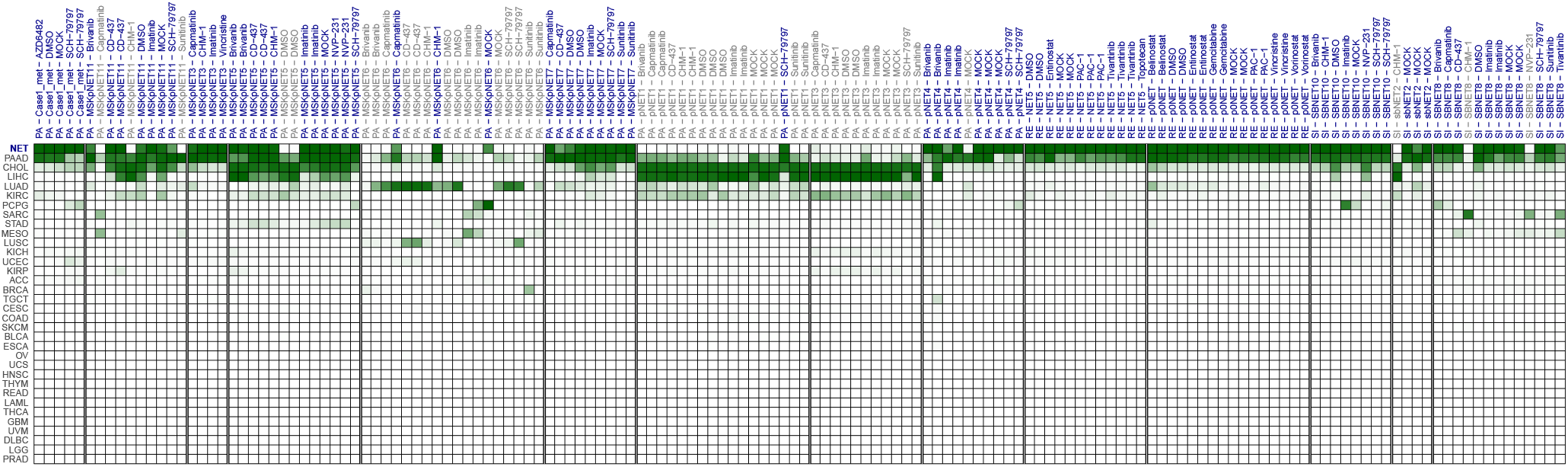
Identification of GEP-NET metastases derived explants expressing GEP-NET specific genes. The heatmap shows the supervised classification likelihood of each explant against 34 tissue types in TCGA. Explant tissue recapitulating the transcriptome of GEP-NET tumors (*S_GEP−NET_ >* 0.5) are highlighted in blue. Other explants are either contaminated by infiltrating tissue or may represent a different tumor histology.

Of 40 tested drugs, 13 were correctly predicted to reverse MR activity (True Positives), 14 were correctly predicted to have no MR reversal activity (True Negatives), 5 were incorrectly predicted to have activity (False Positives), and 8 were incorrectly predicted to have no activity (False Negatives) by H-STS-based OncoTreat analysis (68% prediction accuracy, *p <* 0.05, FET). This shows that, despite their histological differences, H-STS cells represent effective models to predict drug-mediated MR-activity-reversal in explants from *bona fide* GEP-NET patient metastases.

**Fig. 4.**
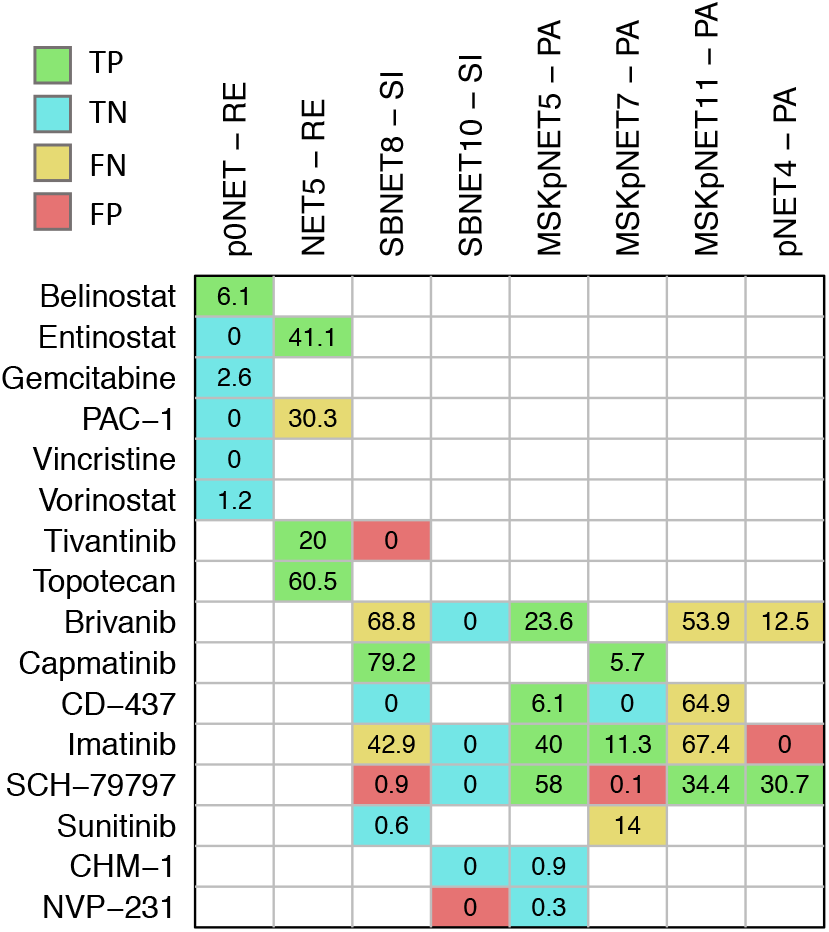
OncoTreat accurately predicts MR-reversing drugs in 8 GEP-NET hepatic metastasis explants, based on H-STS-inferred MoA. Colors indicate the match between H-STS-based OncoTreat-predicted drugs and drugs that experimentally reversed MR-activity in each explant. Specifically, True Positives (TP = 13), True Negatives (TN = 14), False Positives (FP = 5), and False Negatives (FN = 8) are shown in green, cyan, red and yellow, respectively. OncoTreat achieved 68% accuracy, 62% sensitivity, and 74% specificity.

Interestingly, MoA conservation was highly predictive of OncoTreat-predicted drug activity (*p <* 0.001, FET). Indeed, out of 40 drugs evaluated in explant assays, only 3 (7.5%) showed significant MoA conservation but inconsistent OncoTreat-predicted sensitivity (i.e., sensitive when predicted insensitive or vice-versa), and only 4 (10%) showed conservation of predicted OncoTreat activity without statistically significant MoA conservation (Fig. 5).

**Fig. 5.**
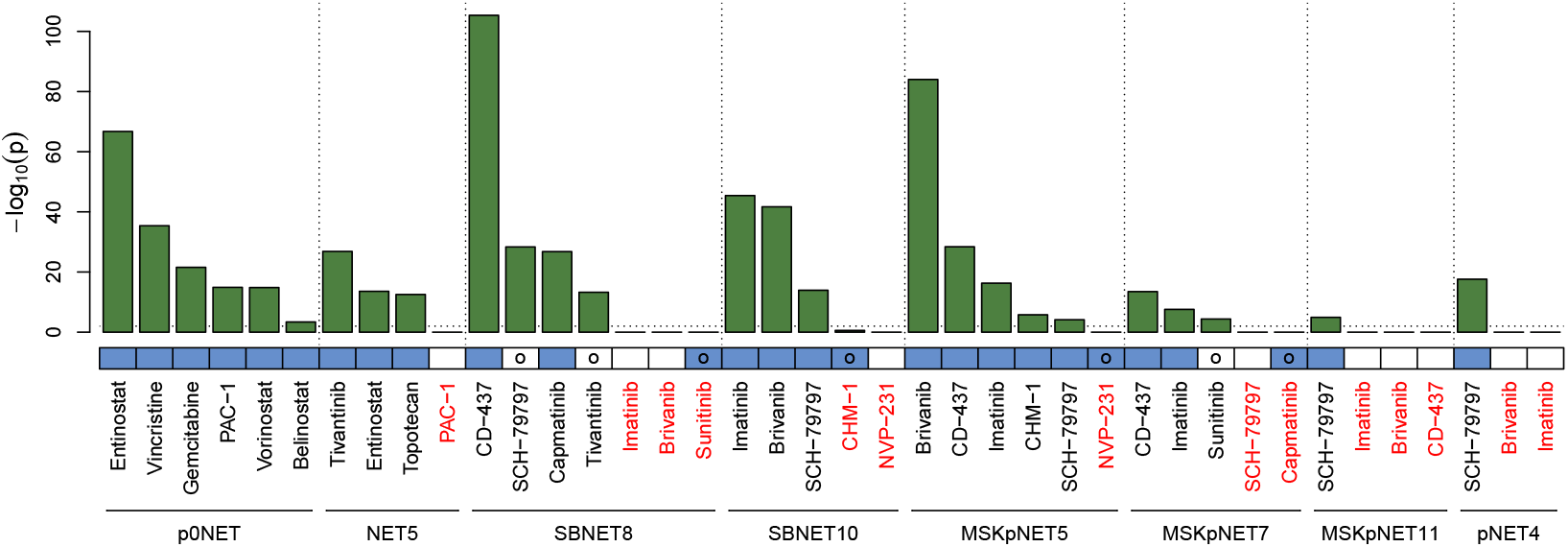
Assessment of drug MoA conservation between GEP-NET metastasis explants and H-STS cells. Bars represent the statistical significance of the MoA conservation, as computed from each explant drug perturbation profile and from the same drug perturbation in H-STS cells. The dotted line represents the statistical threshold to assess MoA conservation (*p <* 0.01, BC). Perturbations showing non-significant MoA conservation are highlighted in red. The blue-colored bar indicates the conservation of H-STS-based and explant-based OncoTreat predictions (*p <* 10^*−*5^, BC). As shown, MoA and OncoTreat overlap are largely consistent, with only 7 exceptions highlighted by circles, including examples with significant MoA conservation but no OncoTreat prediction overlap (SCH-7979 and tivantinib in SBNET8 explant, and sunitinib in MSKpNET7 explant) as well as OncoTreat overlap despite lack of MoA conservation (sunitinib for SBNET8, CHM-1 for SBNET10, NVP-231 for MSKpNET5 and capmatinib for MSKpNET7). All p-values are Bonferroni corrected.

Entinostat was prioritized only for RE-NETs. As such, it was tested only in the two RE-NET-derived explants, where its activity was correctly predicted by H-STS-based On-coTreat. Specifically, consistent with OncoTreat predictions, the analysis showed effective MR-reversal in one explant (NET5, true positive) and correct lack of MR-reversal activity in another (P0NET, true negative) (Fig. 4). Interestingly, also consistent with original results (2), the explant predicted as not entinostat-sensitive was correctly predicted as belinostat-sensitive. Lack of sensitivity to third HDAC-inhibitor, vorinostat, was also correctly predicted, suggesting that H-STS-based OncoTreat analysis is effective at discriminating sensitivity to drugs with closely-related MoA. Consistent with these results, an independent study in *bona fide* NeuroEndocrine Tumor (NET) cells, reported higher sensitivity to HDAC inhibitors compared to control cells (5).

## Discussion

Taken together, these data suggest that while the H-STS and KRJ-1 cells do not represent histology-matched GEP-NET models, they effectively recapitulate (a) drug MoA as assessed in *bona fide* GEP-NET derived cells and (b) critcal regulatory and tumor-dependency features of GEP-NETs. This raises the broader question of what constitutes an optimal model. As discussed in (1), cancer models selected purely based on histology or on the mutational state of key genes can be ineffective in recapitulating patient-relevant tumor biology. Indeed, tumor cells undergo profound reprogramming events, often driven by their aberrant ploidy and epigenetics, yielding transcriptional states that are quite different from their presumed tissue of origin. For instance, another neuroendocrine tumor, Merkel cell carcinoma, was recently shown to be likely B cell rather than Merkel cell derived (11). As a result, the presence of a single genetic alteration of interest, e.g. MYCN amplification, cannot be taken out of context given the large number of additional somatic events —genetic and epigenetic— that contribute to producing immortalized cell lines. We propose that an objective molecular criterion, based on mechanism conservation, as introduced in (2), may provide a valuable additional metric for model selection, even when histology-matched models may not be available.

## ACKNOWLEDGEMENTS

The results in this manuscript were produced using the generous support by the NET Research Foundation to Drs. A.C., D.L.R, and M.H.K for the GEP-NET explant component of the study. We would also like to acknowledge the National Institutes of Health 1R35CA197745 (Outstanding Investigator Award) and U01CA217858 (Cancer Target Discovery and Development network) to A.C., and the NCI instrumentation grants S10OD012351 and S10OD021764 to A.C., which were instrumental for the data analysis. All RNASeq libraries and sequencing was performed in the JP Sulzberger Columbia Genome Center, with support from the P30 Cancer Center Support Grant (P30CA013696).

## Author Contributions

A.C. conceived the study and was involved in all aspects of study organization, sample collection, data analysis and manuscript writing. M.J.A. was responsible for all aspects of data analysis, with help from P.S.M. P.Y., M.L.A., M.B., E.S., C.W.Z., and H.H. were responsible for explant generation, culturing, and treatment. A.G. was responsible for GEP-NET sample preparation, storage and RNA isolation. C.K. and R.R. were responsible for the P0NETCL dissociation, culturing, and treatment to generate drug perturbation profiles. J.A.C., A.T.F., P.E.O., J.W.M., M.H.K., D.L.R. were responsible for patient enrollment into the study. M.L., M.L.A., and G.C. were responsible for the optimization and supervision of the explant generation protocol at the three institutions. D.J. was involved in the study conception and design. Finally, M.J.A. and A.C. were responsible for all aspects of manuscript writing and revision.

### Competing Financial Interests Statement

M.J.A. is Chief Scientific Officer and equity holder at DarwinHealth, Inc., a company that has licensed some of the algorithms used in this manuscript from Columbia University. A.C. is founder, equity holder, consultant and director of DarwinHealth Inc. Columbia University is also an equity holder in DarwinHealth Inc.

### URLs

The VIPER and aREA algorithms are part of the “viper” R-system’s package available from Bioconductor: https://www.bioconductor.org/packages/release/bioc/html/viper.html. The neuroendocrine context-specific regulatory network model is available from Figshare: https://doi.org/10.6084/m9.figshare.6007232. Glioma, pancreas and prostate adenocarcinoma context-specific regulatory networks are available as part of the “aracne.networks” R-system’s package from Bioconductor: https://www.bioconductor.org/packages/release/data/experiment/html/aracne.networks.html.

## Methods

### Cell culture conditions

Neuroendocrine tumor-derived H-STS (7) cells were grown in 1:1 mix of M199:Ham’s F-12 containing 10% FBS and antibiotics (penicillin 100 IU/ml and streptomycin 100 *µ*g/ml). Glioma U87 cell were grown in EMEM containing 10% FBS and antibiotics, HF2597 cells were grown in DMEM/F12 containing 1% N-2 Supplement (Gibco 17502-048), 500 mg/l BSA, 25 mg/l Gentamicin, 0.5% Antibiotic/antimicotic (Invitrogen 15240-062), 20 ng/ml FGF*β* and 20 ng/ml EGF; Pancreas adenocarcinoma AsPC-1 were grown in RPMI-1640 containing 10% FBS and antibiotics, PANC-1 were grown in DMEM supplemented with 10% FBS and antibiotics; Prostate carcinoma LNCaP were grown in phenol red-free RPMI-1640 supplemented with 10% FBS and antibiotics, DU-145 cells were grown in EMEM supplemented with 10% FBS and antibiotics. All cells used in these studies were below passage 30. U87, AsPC-1, PANC-1, LNCaP and DU-145 cells were obtained from ATCC and STR profiled. P0NETCL cells were obtained from a fresh RE-NET hepatic metastasis; necrotic or non-tumor tissue was removed prior to dissociation. The sample was dissociated using the Miltenyi Dissociation Kit, human (130-095-929) using the manufacturer’s protocol. Dissociated cells were filtered through a 70 *µ*m filter and counted and checked for viability using a Countess cell counter (ThermoFisher). P0NETCL cells were grown in RPMI-1640 containing 10% FBS.

### Generation of drug-perturbation databases

For each drug and for each cell line (H-STS, U87, HF2597, AsPC-1, PANC-1, LNCaP and DU-145), the maximum sub-lethal concentration (ED_20_) was established by 10-point dose response curves in triplicate, using total ATP content as read-out. For P0NETCL, drug response curves could not be performed without compromising cell state. Thus, concentration was selected based on the H-STS ED_20_ when assayed in equivalent conditions. Briefly, 2,000 cells/well were plated in 384 well plates. Small molecule compounds were added with a 96 well pin-tool head 12h after cell plating. Viable cells were quantified 48h later by ATP assay (CellTiterGlo, Promega). Relative cell viability was computed using matched DMSO control wells as reference. ED_20_ was estimated by fitting a 4-parameter sigmoid model to the titration results. Then, cells plated in 96 well plates, were perturbed with a library of small molecule compounds at their corresponding ED_20_ concentration and 1/10 of it (Supplementary Table 2). Cells were lysed at 24h after small molecule compound perturbation and sequencing libraries were prepared according to the PLATESeq protocol (10) and sequenced on the HiSeq4000 platform (Illumina) at the Columbia Genome Center. RNA-Seq reads were mapped to the *Homo sapiens* assembly 38 reference genome and counted at the gene level by the STAR aligner software (12). ENSMBL gene identifiers were mapped to Entrez Gene identifiers. The expression data was then normalized by equivariance transformation, based on the negative binomial distribution with the DESeq R-system package (Bioconductor). At least 2 replicates per each condition were obtained.

### Explant treatment prioritization

To optimally preserve cell state, explants cultured in organotypic conditions had to be treated within 24h from collection. As such, drugs could not be prioritized on an explant by explant basis. As a result, we leveraged the top predictions by OncoTreat (2) averaged across each primary tumor organ site to identify a set of drugs for explant treatment (Supplementary Table 1).

### Explant collection, selection, and treatment

Explants were collected, cultured, and treated at three institutions, including Columbia University (IRB:AAAN7562), Dana Farber Cancer Center (IRB:02-314), and Memorial Sloane Kettering (IRB:10-018). Tissues were stored at Columbia Pathology (IRB:AAAB2667). Core needle biopsy specimens, approximately 0.5 cm^3^ to 1.0 cm^3^ in size, were procured from hepatic metastases of GEP-NET patients. Immediately following procurement, touch prep analysis was performed as a Quality Control (QC) measure to determine the presence of viable tumor cells. All specimens passing QC analysis were embedded in non-permanent embedding media (4–6% agarose) and sectioned using a Leica Vibratome VT1000. In general, the softer the tissue the slower was the slicing speed (0.03–0.08 mm/sec neoplastic tissue; 0.01–0.08 mm/sec normal tissue). Vibration amplitude was set at 2.95–3.0 mm. Precision-sliced sections 200–400 *µ*m thick (minus permanent embedding media) were transferred immediately to a multi-well plate containing media and allowed to acclimate for one hour prior to drug exposure. Optimal drug concentrations were determined according to ED_20_ data in cell lines, see Supplementary Table 1. All treatments were performed in duplicate, under standard tissue culture conditions. Treated sections were OCT-embedded. From the fresh frozen sample, a section was H&E stained to confirm presence and abundance of tumor cells. Once confirmed total RNA was extracted for RNASeq profiling, using the TruSeq library preparation protocol and sequenced on an HiSeq 4000 (Illumina) at the Columbia Genome Center. RNASeq reads were mapped to the *Homo sapiens* assembly 38 reference genome and counted at the gene level by the STAR aligner software (12). ENSMBL gene identifiers were mapped to Entrez Gene identifiers. The expression data was then normalized by equivariance transformation, based on the negative binomial distribution with the DESeq R-system package (Bioconductor).

### Explants quality assessment

In total, 150 samples from 14 GEP-NET hepatic metastases were analyzed. We used supervised machine learning to identify explants as *bona fide* GEP-NET derived, using a GEP-NET classifier. Specifically, for the positive and negative controls, the gene expression probability density of each gene was estimated by fitting a 2-gaussian mixture model to the log-transformed, RPKM-normalized expression of either 212 patient-derived GEP-NET profiles (2) or 10,900 tumor samples in TCGA, representing 33 distinct tumor types in total. A set of 1,600 genes that optimally discriminate each tumor type, including GEP-NET, against the background of all other tumors was assembled by selecting the top 100 genes most differentially expressed in each of the 33 tumor types. Then, we quantified the gene expression profile similarity of each explant sample against each GEP-NET and TCGA samples by correlation analysis. Thus, each explant *E*_*i*_ was associated with a ranked vector *X*_*i*_, representing the similarity of the explant to each of the 11,112 reference tumor samples. Finally, a tumor type score was computed for each explant, based on the enrichment of the vector *X*_*i*_ in samples from that tumor type, by gene set enrichment analysis, using the aREA algorithm (4). Scores were scaled based on the maximum possible enrichment score for each tumor-type. The relative likelihood for the GEP-NET class (SGEP-NET) was computed from the distribution of GEP-NET scores for *bona fide* GEP-NET tumors (212 samples) vs. the corresponding scores for all the non GEP-NET tumors (10,900 samples). Of the 150 drug-treated and control explant samples, 99 showed a GEP-NET tumor-type score SGEP-NET *>* 0.5, and were considered as *bona fide* GEP-NET-derived (Supplementary Fig. 2). Finally, we removed explants with low tissue availability, which were perturbed with fewer than 3 drugs, generating a final list comprising 83 high-quality samples from 8 distinct explants, including 2 RE-NETs: P0NET and NET5; 2 SI-NETs: SBNET8 and SBNET10, and 4 P-NETs: MSKpNET5, MSKpNET7, MSKpNET11 and pNET4.

### MoA Analysis

Differential gene expression signatures were computed for H-STS and P0NETCL by comparing profiles from each drug-treated sample with those from plate-matched vehicle controls (DMSO), using a moderated Student’s t-test as implemented in the limma package from Bioconductor (13). Gene expression signatures for the explants were computed by comparing each drug perturbation vs. the explant-matched vehicle control, with exception of MSKpNET5 and pNET4 that, due to QC limitations (GEP-NET transcriptome recapitulation), were compared against untreated controls. Individual gene expression signatures were then transformed into protein activity signatures, based on the GEP-NET regulatory network —with exception of the positive controls shown in Fig. 2a, for which tissue lineage-matched regulatory networks were used— using the VIPER algorithm (4), as implemented in the viper package from Bioconductor.

### Comparative MoA analysis in P0NETCL cells

MoA similarity between P0NETCL and other cell lines was quantified by comparing drug-induced protein activity signatures. For this, a reciprocal enrichment analysis was performed (14) by computing the enrichment of the top/bottom 50 most differentially active proteins in response to drug perturbation in cell line “A” on the drug-induced protein activity signature of cell line “B”, and then by integrating (average) the result (normalized enrichment score) with the corresponding one to the enrichment of the top/bottom 50 most differentially active proteins in cell line “B” on the drug-induced protein activity of cell line “A”. Enrichment and p-value were computed by the aREA algorithm (4). When two concentrations for a given drug were available, the concentration showing the strongest MoA conservation was used. Drug perturbation bioactivity was computed as the area over the cumulative distribution for the absolute value of drug-induced differential protein activity, expressed as the normalized enrichment score computed by the aREA algorithm (9), and scaled to the maximum value across all drug perturbations.

To highlight the relationship between cell-line-specific drug bioactivity and MoA conservation, for each cell line pair to be compared, we rank-sorted all drug perturbations based on the lower of the two bioactivities assessed in each cell line pair. Then, we averaged over a running-window of size 40-drugs to compute the fraction of drugs showing significant MoA conservation (*p <* 10^*−*10^, BC). The 40-drug window whose average bioactivity was closest to the entinostat bioactivity was highlighted with a red circle (Fig. 2a).

Drug MoA conservation was also evaluated for the perturbations performed in the GEP-NET patient-derived explants, by comparing their MoA to the corresponding one from the matching drug perturbations performed in H-STS cells.

### OncoTreat Analysis

OncoTreat analysis was performed as previously described (2). Briefly, drugs were prioritized for each GEP-NET hepatic metastasis sample and explant based on their ability to invert their master regulator program. As previously described (2), gene expression signatures were obtained for each GEP-NET hepatic metastasis sample by comparison against the pool of cluster-matched primary samples. For the explants, gene expression signatures were obtained by comparing the transcriptome of each explant against the tissue of origin-matched primary tumors. Protein activity signatures were subsequently derived from the gene expression signatures by VIPER analysis (4). Master regulator proteins were identified as the top/bottom 50 most differentially active proteins in each GEP-NET hepatic metastasis tumor and explant. Finally, drugs were prioritized by computing the enrichment of the master regulator proteins of each explant on each drug-induced protein activity signature with the aREA algorithm (4). P-values were estimated by the analytical approximation implemented in the aREA algorithm, which have been shown to be practically equivalent to estimations obtained by permuting the proteins in the signature uniformly at random (4). P-values were corrected to account for multiple hypothesis testing by the Bonferroni’s method.

### Comparative OncoTreat analysis in P0NETCL cells

Overlap of either H-STS-based or P0NETCL-based OncoTreat drug predictions for 69 GEP-NET hepatic metastases was assessed and compared to the overlap of H-STS-based OncoTreat inferences with those based on perturbational profiles from 6 additional unrelated cell lines (negative controls). Drugs were rank-sorted based on their bioactivity, using the minimum bioactivity computed in each pair of selected cells. Results were averaged over a running-window of 40 drugs to increase the number of data points used to compute the overlap between OncoTreat predictions and also to smooth the results as a function of drug bioactivity. The overlap of drugs considered as significantly inverting the pattern of activity of GEP-NET hepatic metastases (*p <* 10^*−*5^, BC) was quantified by the odd’s ratio and statistical significance estimated by the Fisher’s exact test.

### Statistical analysis

Enrichment analysis, including model matching based on MR conservation, MoA conservation, and OncoTreat analysis, was computed by the aREA algorithm and statistical significance was estimated by the analytical approximation implemented in the algorithm (4). The significance for the overlap of OncoTreat predictions was estimated by the Fisher’s exact test. Statistical significance for the difference in odds’ ratio when comparing the conservation of OncoTreat results was estimated by one-sample Wilcoxon’s signed rank test (U-test). All statistical tests were performed with the R-system. All p-values were Bonferroni corrected to account for multiple hypothesis testing.

### Code availability

All the code used in this work is freely available for research purposes. VIPER and aREA algorithms are part of the “viper” R-system’s package available from Bioconductor (see URLs). The context-specific regulatory network models are available from Figshare and Bioconductor (see URLs).

---

**Supplementary Table 1.**
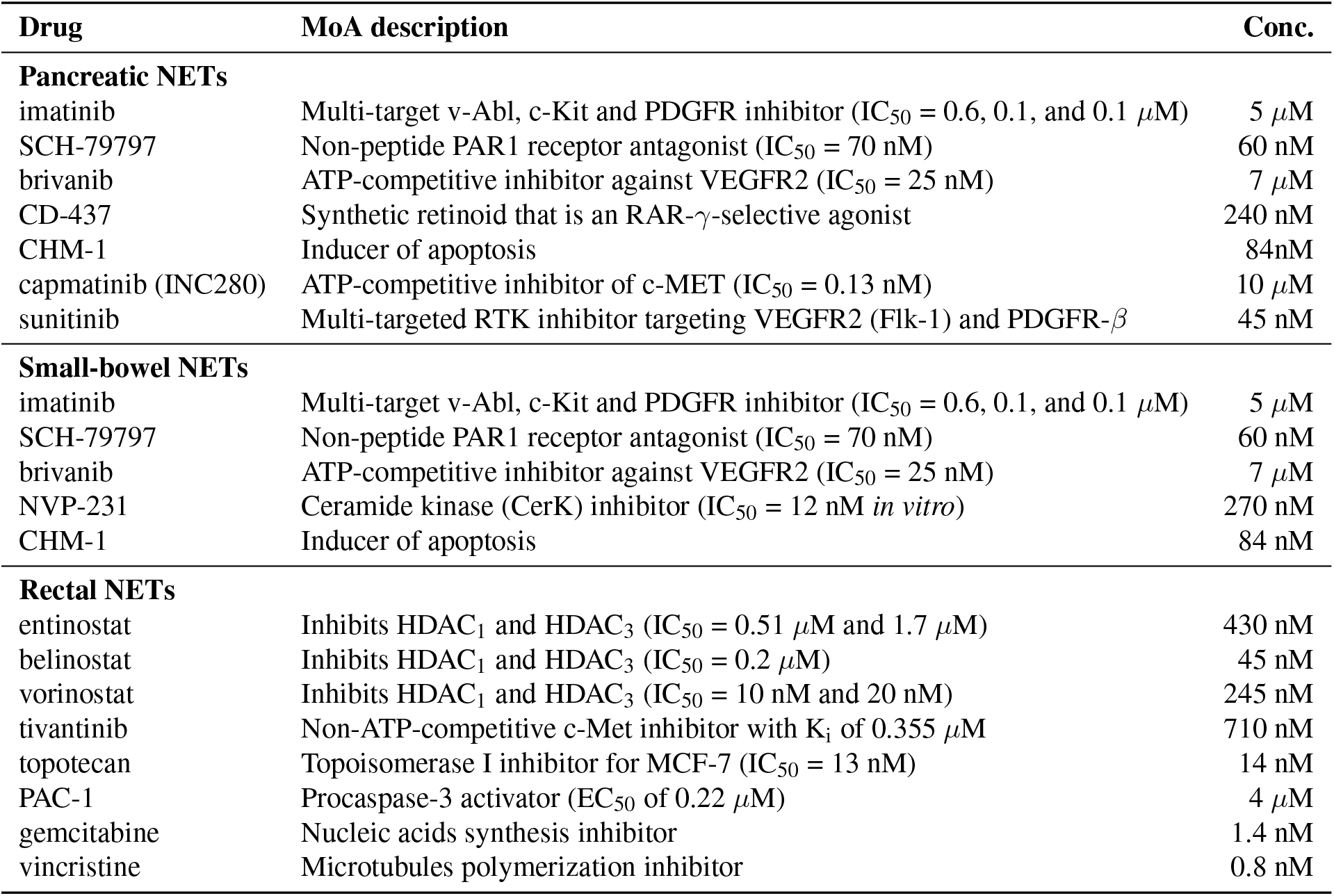
Drugs prioritized for the GEP-NET hepatic metastasis explants perturbations. The table shows a short MoA description and the concentration used, which was estimated as the ED_20_ at 48h in dose-response cell viability assays.

| Drug | MoA description | Conc. |
| --- | --- | --- |
| **Pancreatic NETs** |  |  |
| imatinib | Multi-target v-Abl, c-Kit and PDGFR inhibitor (IC<sub>50</sub> = 0.6, 0.1, and 0.1 μM) | 5 μM |
| SCH-79797 | Non-peptide PAR1 receptor antagonist (IC<sub>50</sub> = 70 nM) | 60 nM |
| brivanib | ATP-competitive inhibitor against VEGFR2 (IC<sub>50</sub> = 25 nM) | 7 μM |
| CD-437 | Synthetic retinoid that is an RAR-γ-selective agonist | 240 nM |
| CHM-1 | Inducer of apoptosis | 84nM |
| capmatinib (INC280) | ATP-competitive inhibitor of c-MET (IC<sub>50</sub> = 0.13 nM) | 10 μM |
| sunitinib | Multi-targeted RTK inhibitor targeting VEGFR2 (Flk-1) and PDGFR-β | 45 nM |
| **Small-bowel NETs** |  |  |
| imatinib | Multi-target v-Abl, c-Kit and PDGFR inhibitor (IC<sub>50</sub> = 0.6, 0.1, and 0.1 μM) | 5 μM |
| SCH-79797 | Non-peptide PAR1 receptor antagonist (IC<sub>50</sub> = 70 nM) | 60 nM |
| brivanib | ATP-competitive inhibitor against VEGFR2 (IC<sub>50</sub> = 25 nM) | 7 μM |
| NVP-231 | Ceramide kinase (CerK) inhibitor (IC<sub>50</sub> = 12 nM *in vitro*) | 270 nM |
| CHM-1 | Inducer of apoptosis | 84 nM |
| **Rectal NETs** |  |  |
| entinostat | Inhibits HDAC<sub>1</sub> and HDAC<sub>3</sub> (IC<sub>50</sub> = 0.51 μM and 1.7 μM) | 430 nM |
| belinostat | Inhibits HDAC<sub>1</sub> and HDAC<sub>3</sub> (IC<sub>50</sub> = 0.2 μM) | 45 nM |
| vorinostat | Inhibits HDAC<sub>1</sub> and HDAC<sub>3</sub> (IC<sub>50</sub> = 10 nM and 20 nM) | 245 nM |
| tivantinib | Non-ATP-competitive c-Met inhibitor with K<sub>i</sub> of 0.355 μM | 710 nM |
| topotecan | Topoisomerase I inhibitor for MCF-7 (IC<sub>50</sub> = 13 nM) | 14 nM |
| PAC-1 | Procaspase-3 activator (EC<sub>50</sub> of 0.22 μM) | 4 μM |
| gemcitabine | Nucleic acids synthesis inhibitor | 1.4 nM |
| vincristine | Microtubules polymerization inhibitor | 0.8 nM |

**Supplementary Table 2.**
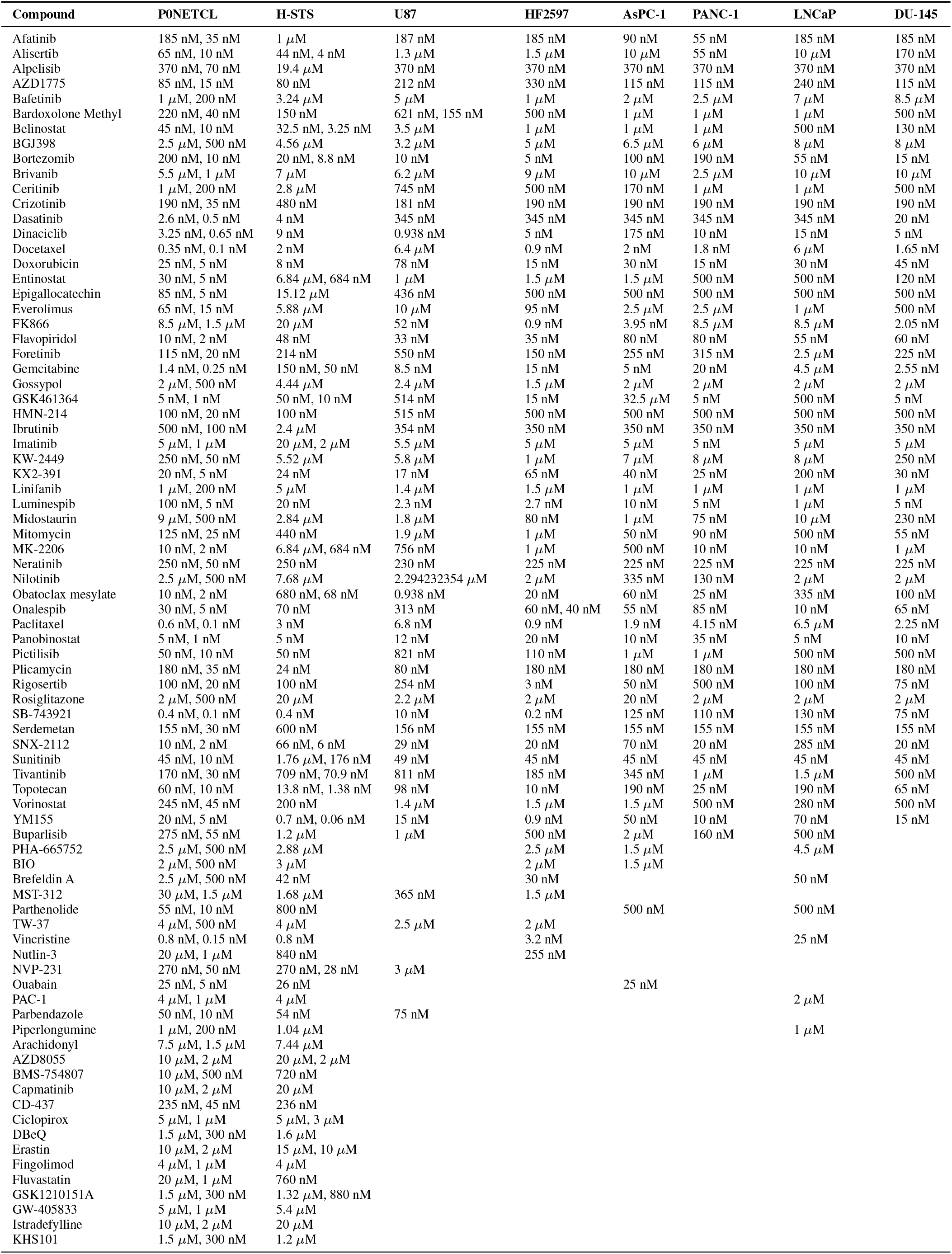
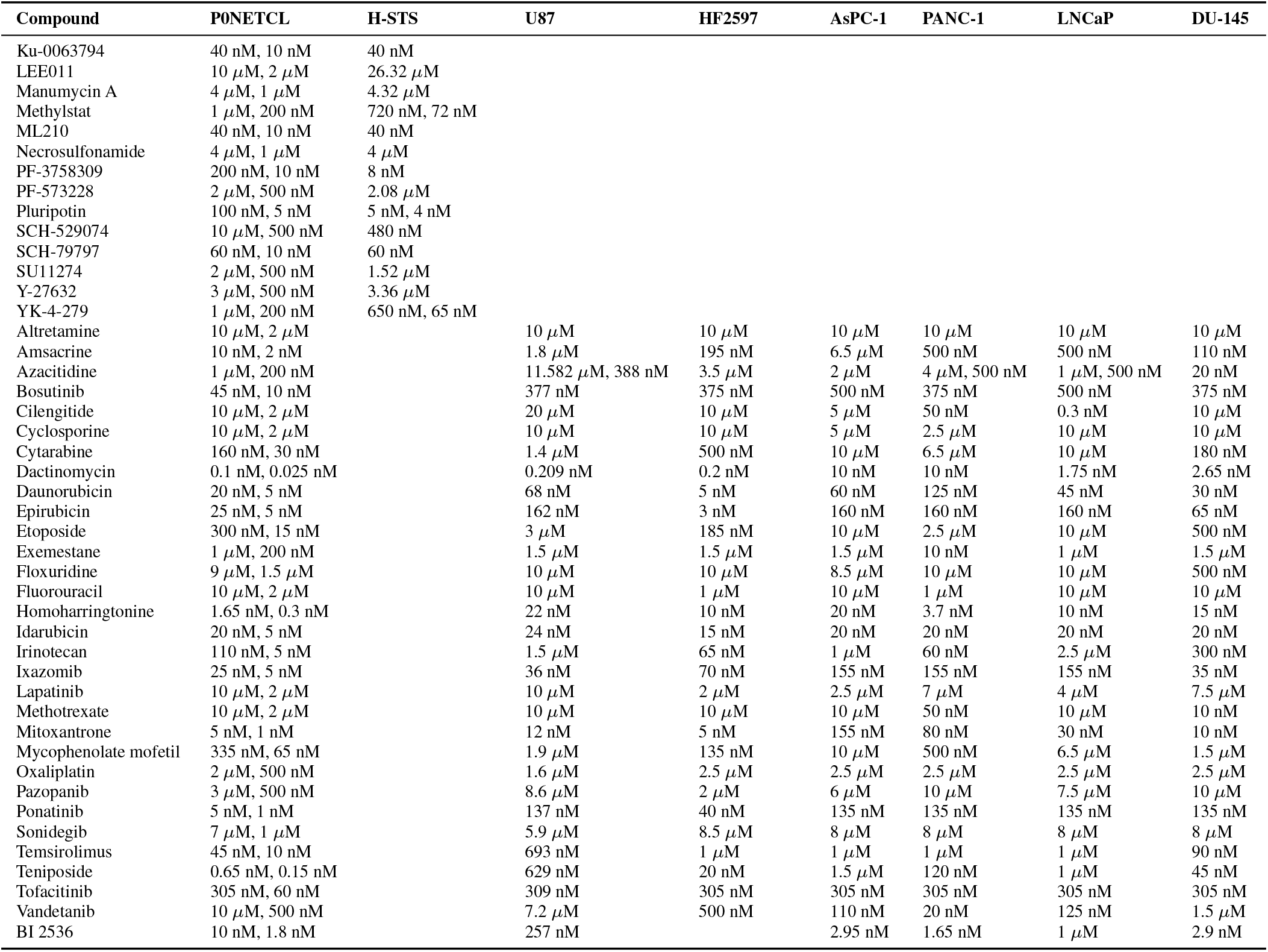
Drug concentrations used to perturb 8 cell lines.

## Bibliography

1. P. Rajbhandari, G. Lopez, C. Capdevila, B. Salvatori, J. Y. Yu, R. Rodriguez-Barrueco, D. Martinez, M. Yarmarkovich, N. Weichert-Leahey, B. J. Abraham, M. J. Alvarez, A. Iyer, J. L. Harenza, D. Oldridge, K. De Preter, J. Koster, S. Asgharzadeh, R. C. Seeger, J. S. Wei, J. Khan, J. Vandesompele, P. Mestdagh, R. Versteeg, A. T. Look, R. A. Young, A. Iavarone, A. Lasorella, J. M. Silva, J. M. Maris, and A. Califano. Cross-cohort analysis identifies a tead4-mycn positive feedback loop as the core regulatory element of high-risk neuroblastoma. Cancer Discovery, 8(5):582–599, 2018. ISSN 2159-8274. doi: 10.1158/2159-8290.Cd-16-0861.

2. M. J. Alvarez, P. S. Subramaniam, L. H. Tang, A. Grunn, M. Aburi, G. Rieckhof, E. V. Komissarova, E. A. Hagan, L. Bodei, P. A. Clemons, F. S. Dela Cruz, D. Dhall, D. Diolaiti, D. A. Fraker, A. Ghavami, D. Kaemmerer, C. Karan, M. Kidd, K. M. Kim, H. C. Kim, L. P. Kunju, U. Langel, Z. Li, J. Lee, H. Li, V. LiVolsi, R. Pfragner, A. R. Rainey, R. B. Realubit, H. Remotti, J. Regberg, R. Roses, A. Rustgi, A. R. Sepulveda, S. Serra, C. Shi, X. Yuan, M. Barberis, R. Bergamaschi, A. M. Chinnaiyan, T. Detre, S. Ezzat, A. Frilling, M. Hommann, D. Jaeger, M. K. Kim, B. S. Knudsen, A. L. Kung, E. Leahy, D. C. Metz, J. W. Milsom, Y. S. Park, D. Reidy-Lagunes, S. Schreiber, K. Washington, B. Wiedenmann, I. Modlin, and A. Califano. A precision oncology approach to the pharmacological targeting of mechanistic dependencies in neuroendocrine tumors. Nat Genet, 50(7):979–989, 2018. ISSN 1546-1718 (Electronic) 1061-4036 (Linking). doi: 10.1038/s41588-018-0138-4.

3. A. Califano and M. J. Alvarez. The recurrent architecture of tumour initiation, progression and drug sensitivity. Nat Rev Cancer, 17(2):116–130, 2017. ISSN 1474-1768 (Electronic) 1474-175X (Linking). doi: 10.1038/nrc.2016.124.

4. M. J. Alvarez, Y. Shen, F. M. Giorgi, A. Lachmann, B. B. Ding, B. H. Ye, and A. Califano. Functional characterization of somatic mutations in cancer using network-based inference of protein activity. Nat Genet, 48(8):838–47, 2016. ISSN 1546-1718 (Electronic) 1061-4036 (Linking). doi: 10.1038/ng.3593.

5. T. Hofving, Y. Arvidsson, B. Almobarak, L. Inge, R. Pfragner, M. Persson, G. Stenman, E. Kristiansson, V. Johanson, and O. Nilsson. The neuroendocrine phenotype, genomic profile and therapeutic sensitivity of gepnet cell lines. Endocr Relat Cancer, 25(4):X1–X2, 2018. ISSN 1479-6821 (Electronic) 1351-0088 (Linking). doi: 10.1530/ERC-17-0445e.

6. J. Barretina, G. Caponigro, N. Stransky, K. Venkatesan, A. A. Margolin, S. Kim, C. J. Wilson, J. Lehar, G. V. Kryukov, D. Sonkin, A. Reddy, M. Liu, L. Murray, M. F. Berger, J. E. Monahan, P. Morais, J. Meltzer, A. Korejwa, J. Jane-Valbuena, F. A. Mapa, J. Thibault, E. Bric-Furlong, P. Raman, A. Shipway, I. H. Engels, J. Cheng, G. K. Yu, J. Yu, Jr. Aspesi, P., M. de Silva, K. Jagtap, M. D. Jones, L. Wang, C. Hatton, E. Palescandolo, S. Gupta, S. Mahan, C. Sougnez, R. C. Onofrio, T. Liefeld, L. MacConaill, W. Winckler, M. Reich, N. Li, J. P. Mesirov, S. B. Gabriel, G. Getz, K. Ardlie, V. Chan, V. E. Myer, B. L. Weber, J. Porter, M. Warmuth, P. Finan, J. L. Harris, M. Meyerson, T. R. Golub, M. P. Morrissey, W. R. Sellers, R. Schlegel, and L. A. Garraway. The cancer cell line encyclopedia enables predictive modelling of anticancer drug sensitivity. Nature, 483(7391):603–7, 2012. ISSN 1476-4687 (Electronic) 0028-0836 (Linking). doi: 10.1038/nature11003.

7. R. Pfragner, A. Behmel, H. Hoger, A. Beham, E. Ingolic, I. Stelzer, B. Svejda, V. A. Moser, A. C. Obenauf, V. Siegl, O. Haas, and B. Niederle. Establishment and characterization of three novel cell lines-p-sts, l-sts, h-sts - derived from a human metastatic midgut carcinoid. Anticancer Res, 29(6):1951–61, 2009. ISSN 0250-7005 (Print) 0250-7005 (Linking).

8. B. Rinner, B. Galle, S. Trajanoski, C. Fischer, M. Hatz, T. Maierhofer, G. Michelitsch, F. Moinfar, I. Stelzer, R. Pfragner, and C. Guelly. Molecular evidence for the bi-clonal origin of neuroendocrine tumor derived metastases. BMC Genomics, 13:594, 2012. ISSN 1471-2164 (Electronic) 1471-2164 (Linking). doi: 10.1186/1471-2164-13-594.

9. Y. Shen, M. J. Alvarez, B. Bisikirska, A. Lachmann, R. Realubit, S. Pampou, J. Coku, C. Karan, and A. Califano. Systematic, network-based characterization of therapeutic target inhibitors. PLoS Comput Biol, 13(10):e1005599, 2017. ISSN 1553-7358 (Electronic) 1553-734X (Linking). doi: 10.1371/journal.pcbi.1005599.

10. E. C. Bush, F. Ray, M. J. Alvarez, R. Realubit, H. Li, C. Karan, A. Califano, and P. A. Sims. Plate-seq for genome-wide regulatory network analysis of high-throughput screens. Nat Commun, 8(1):105, 2017. ISSN 2041-1723 (Electronic) 2041-1723 (Linking). doi: 10.1038/s41467-017-00136-z.

11. C. M. Sauer, A. M. Haugg, E. Chteinberg, D. Rennspiess, V. Winnepenninckx, E. J. Speel, J. C. Becker, A. K. Kurz, and A. Zur Hausen. Reviewing the current evidence supporting early b-cells as the cellular origin of merkel cell carcinoma. Crit Rev Oncol Hematol, 116: 99–105, 2017. ISSN 1879-0461 (Electronic) 1040-8428 (Linking). doi: 10.1016/j.critrevonc. 2017.05.009.

12. A. Dobin, C. A. Davis, F. Schlesinger, J. Drenkow, C. Zaleski, S. Jha, P. Batut, M. Chaisson, and T. R. Gingeras. Star: ultrafast universal rna-seq aligner. Bioinformatics, 29(1):15–21, 2013. ISSN 1367-4811 (Electronic) 1367-4803 (Linking). doi: 10.1093/bioinformatics/bts635.

13. M. E. Ritchie, B. Phipson, D. Wu, Y. Hu, C. W. Law, W. Shi, and G. K. Smyth. limma powers differential expression analyses for rna-sequencing and microarray studies. Nucleic Acids Res, 43(7):e47, 2015. ISSN 1362-4962 (Electronic) 0305-1048 (Linking). doi: 10.1093/nar/gkv007.

14. M. Kruithof-de Julio, M. J. Alvarez, A. Galli, J. Chu, S. M. Price, A. Califano, and M. M. Shen. Regulation of extra-embryonic endoderm stem cell differentiation by nodal and cripto signaling. Development, 138(18):3885–95, 2011. ISSN 1477-9129 (Electronic) 0950-1991 (Linking). doi: 10.1242/dev.065656.

